# Prenatal exposure to valproic acid reduces synaptic δ-catenin levels and disrupts ultrasonic vocalization in neonates

**DOI:** 10.1101/2023.12.14.571709

**Authors:** Seung Hyun Roh, Hadassah Mendez-Vazquez, Matheus F. Sathler, Michael J. Doolittle, Anastasiya Zaytseva, Hannah Brown, Morgan Sainsbury, Seonil Kim

## Abstract

Valproic acid (VPA) is an effective and commonly prescribed drug for epilepsy and bipolar disorder. However, children born from mothers treated with VPA during pregnancy exhibit an increased incidence of autism spectrum disorder (ASD). Although VPA may impair brain development at the cellular level, the mechanism of VPA-induced ASD has not been completely addressed. A previous study has found that VPA treatment strongly reduces δ-catenin mRNA levels in cultured human neurons. δ-catenin is important for the control of glutamatergic synapses and is strongly associated with ASD. VPA inhibits dendritic morphogenesis in developing neurons, an effect that is also found in neurons lacking δ-catenin expression. We thus hypothesize that prenatal exposure to VPA significantly reduces δ-catenin levels in the brain, which impairs glutamatergic synapses to cause ASD. Here, we found that prenatal exposure to VPA markedly reduced δ-catenin levels in the brain of mouse pups. VPA treatment also impaired dendritic branching in developing mouse cortical neurons, which was reversed by elevating δ-catenin expression. Prenatal VPA exposure significantly reduced synaptic AMPA receptor levels and postsynaptic density 95 (PSD95) in the brain of mouse pups, indicating dysfunctions in glutamatergic synaptic transmission. VPA exposure also significantly altered ultrasonic vocalization (USV) in newly born pups when they were isolated from their nest. Moreover, VPA-exposed pups show impaired hypothalamic response to isolation, which is required to produce animals’ USVs following isolation from the nest. Therefore, these results suggest that VPA-induced ASD pathology can be mediated by the loss of δ-catenin functions.

**Highlights:**

- Prenatal exposure of valproic acid (VPA) in mice significantly reduces synaptic δ-catenin protein and AMPA receptor levels in the pups’ brains.
- VPA treatment significantly impairs dendritic branching in cultured cortical neurons, which is reversed by increased δ-catenin expression.
- VPA exposed pups exhibit impaired communication such as ultrasonic vocalization.
- Neuronal activation linked to ultrasonic vocalization is absent in VPA-exposed pups.
- The loss of δ-catenin functions underlies VPA-induced autism spectrum disorder (ASD) in early childhood.

## 1. Introduction

Autism spectrum disorder (ASD) is one of the most heterogeneous neurodevelopmental disorders, but there is no known single cause for ASD (Sandin et al., 2017; Yoo, 2015). Although there is a significant genetic component to ASD, studies suggest that environmental factors may also play a role in the development of the disorder (Christensen et al., 2013; Huguet et al., 2013). Several environmental factors have been investigated for their potential association with ASD, including prenatal and early-life exposures (Karimi et al., 2017). For example, maternal infections during pregnancy, such as rubella or cytomegalovirus, have been linked to an increased risk of ASD (Deykin & MacMahon, 1979; Eidelman & Samueloff, 2002; Libbey et al., 2005; Ornoy et al., 2015; Taylor et al., 2002). Moreover, maternal exposure to certain medications, such as valproic acid or thalidomide, has also been associated with a higher likelihood of ASD in some studies (Ito et al., 2010; Narita et al., 2010; Ornoy et al., 2015). Additional studies have further explored the role of environmental toxins, such as air pollutants, heavy metals (e.g., lead, mercury), pesticides, and certain chemicals, in autism development (Rossignol et al., 2014).

Exposure to exogenous drugs during pregnancy may interfere with brain development and maturation in the fetus; one such exogenous chemical is valproic acid (VPA). VPA is an effective and commonly prescribed drug for epilepsy and bipolar disorder. In the early 1980s, however, VPA was found to be teratogenic (Brown et al., 1980; Robert & Guibaud, 1982). Children born to mothers who were treated with VPA during pregnancy exhibit an increased incidence of neurological diseases, including ASD (Jentink et al., 2010; Koren et al., 2006; Moore et al., 2000; Rasalam et al., 2005; Roullet et al., 2013; Williams et al., 2001). Although VPA may impair brain development at the cellular level, the mechanism of VPA’s teratogenic effects are not well characterized. A recent study using human embryonic/induced pluripotent stem (ES/iPS) cell-derived glutamatergic neurons has found that VPA treatment significantly reduces δ-catenin mRNA levels (Chanda et al., 2019). Several human genetic studies suggest that the δ*-catenin* gene is strongly linked to ASD (Guo et al., 2018; Miller et al., 2019; Tuncay et al., 2022; Turner et al., 2015; Wang et al., 2016). The δ*-catenin* gene harbors many deleterious missense mutations, copy number variation, and *de novo* mutations that are strongly associated with severely affected ASD patients from multiple families (Turner et al., 2015; Wang et al., 2016). Studies, including our own, demonstrate that a loss of δ-catenin functions contributes to the ASD-linked cellular pathology and social dysfunction (Mendez-Vazquez et al., 2023; Turner et al., 2015). Therefore, these results suggest that VPA-induced ASD can be mediated by the loss of δ-catenin functions.

Although ASD is one of the most heterogeneous neurodevelopmental disorders (Sandin et al., 2017; Yoo, 2015), many of the ASD-risk genes are directly involved in synapse formation and function (Betancur et al., 2009; Bourgeron, 2015; Chung et al., 2012; El-Amraoui & Petit, 2010; Hampson et al., 2012; Lepeta et al., 2016). δ-catenin is a member of the armadillo repeat protein family, and is highly expressed in neurons and specifically localizes to the excitatory post-synapses (Gilbert & Man, 2016; Peifer et al., 1994; Silverman et al., 2007; Takeichi, 1988). At the postsynaptic density (PSD), δ-catenin interacts with the intracellular domain of N-cadherin, a synaptic cell adhesion protein, and the carboxyl-terminus of δ-catenin binds to AMPA receptor-binding protein (ABP) and glutamate receptor-interacting protein (GRIP) (Gilbert & Man, 2016; Kosik et al., 2005; Silverman et al., 2007; Yuan et al., 2015). This N-cadherin-δ-catenin-ABP/GRIP complex functions as an anchorage for the AMPA-type glutamate receptor (AMPAR) subunit GluA2 (Silverman et al., 2007) and plays a crucial role in maintaining synaptic function and dendrite structure. VPA inhibits dendritic morphogenesis in developing human and rodent neurons, resulting in impaired neurite arborization, reduction of dendrite length and branching (Baumert et al., 2020; Chanda et al., 2019; Matter et al., 2009), which is also evident in neurons lacking δ-catenin expression (Baumert et al., 2020; Israely et al., 2004; Matter et al., 2009; Restituito et al., 2011). In addition to structural deficits, VPA exposure decreases synaptic strength, as measured by AMPAR-mediated excitatory postsynaptic current (EPSC) amplitude (Chanda et al., 2019). Research, including our own, reveals that δ-catenin deficiency increases the synaptic excitation and inhibition ratio and intrinsic excitability in young cortical pyramidal neurons (Assendorp et al., 2022; Mendez-Vazquez et al., 2023). Additionally, both short-term and long-term synaptic plasticity are significantly impaired in the hippocampus of δ-catenin knockout (KO) mice (Israely et al., 2004). This suggests that VPA-induced ASD pathology may result from the loss of δ-catenin function-mediated defects in synaptic properties.

Here, we discover that VPA disrupts the development of cultured mouse cortical neurons, which is reversed by elevating δ-catenin expression. Prenatal exposure to VPA significantly decreases synaptic δ-catenin and AMPAR levels. This likely contributes to abnormalities in hypothalamic neuronal activity, which underlies impaired ultrasonic vocalization (USV) in newly born pups. Therefore, these results suggest that VPA-induced ASD pathology can be mediated by the loss of δ-catenin functions.

## 2. Material and Methods

### 2.1. Animals

CD-1 mice were obtained from Charles River (022) and bred in the animal facility at Colorado State University (CSU). Animals were housed under a 12:12 hour light/dark cycle. CSU’s Institutional Animal Care and Use Committee (IACUC) reviewed and approved the animal care and protocol (3408).

### 2.2. *In vivo* VPA treatment

VPA administration as a single dose (250-500 mg/kg) to pregnant female mice at a critical stage of embryonic development (E12.5) has been consistently shown to be a reliable ASD model recapitulating several behavioral, cellular and molecular phenotypes of the disorder (Al Sagheer et al., 2018; Chaliha et al., 2020; Kataoka et al., 2013; Nicolini & Fahnestock, 2018; Rodier et al., 1996; Schneider & Przewlocki, 2005; Thabault et al., 2022). Therefore, the E12.5 pregnant animals received a single intraperitoneal (IP) injection of 250 mg/kg VPA dissolved in saline, or an equal volume of saline as a control (CTRL). We observed an immediate reduction in animal activity within 5-10 min of VPA injection, consistent with the previous finding (Chanda et al., 2019). Pups were sacrificed on postnatal day 10 (P10), and brain tissues were collected for biochemical analysis.

### 2.3. Primary cortical neuronal culture

Postnatal day 0 (P0) male and female CD-1 pups were used to produce mouse cortical neuron cultures as shown previously (Mendez-Vazquez et al., 2023; Sathler et al., 2022; Sathler et al., 2021; Sztukowski et al., 2018). Cortices were isolated from P0 CD-1 mouse brain tissues and digested with 10 U/mL papain (Worthington Biochemical Corp., LK003176). Mouse cortical neurons were plated on following poly lysine-coated glass bottom dishes (500,000 cells). Neurons were grown in Neurobasal Medium without phenol red (Thermo Fisher Scientific, 12348017) with B27 supplement (Thermo Fisher Scientific, 17504044), 0.5 mM Glutamax (Thermo Fisher Scientific, 35050061), and 1% penicillin/streptomycin (Thermo Fisher Scientific, 15070063).

### 2.4. Dendritic morphology analysis

To visualize dendritic morphology, primary mouse cortical neurons were transfected with pEGFP-C1 vector (Clontech) at 3 days *in vitro* (DIV) using Lipofectamine 2000 (Life Technologies) according to the manufacturer’s protocol as shown previously (Kim & Ziff, 2014; Roberts et al., 2021; Sun et al., 2019; Sztukowski et al., 2018). To express δ-catenin, neurons were transfected with pcDNA3.1-δ-catenin (Mendez-Vazquez et al., 2023) at 3 DIV using Lipofectamine 2000 as described above. 1 mM VPA or saline was treated in neurons at 4 DIV for 72 hours, and cells were fixed at 7 DIV, a condition that is optimal to evaluate neuronal development and dendrite morphology in cultured neurons (Baumert et al., 2020). Neurons were visualized using the Olympus inverted microscope IX73 with an Apo-Plan 20x objective. Images were captured as z-series at a step size of 1 μm with the Olympus cellSENS Dimensions software. For Sholl analysis, maximum-projection images were converted to binary and subjected to analysis in the NIH ImageJ software (https://imagej.nih.gov/ij/). Concentric rings with a step size of 20 µm were used to quantify numbers of dendrite branching as it relates to distance from the soma.

### 2.5. USV Recording and analysis

The ages of the pups used for experiments ranged from postnatal day 5 to postnatal day 7 (P5-P7) as shown previously (Tran et al., 2021). Neonatal P5-7 mice were separated from the home cage and placed in a round transparent container with an open top. The ultrasound microphone (Avisoft-Bioacoustics CM16/CMPA) was placed above the container in the sound-proof box. Vocalizations were recorded for 5 min using Avisoft Recorder software. Between animals, the container was cleaned with 70% ethanol and dried to get rid of any odors that could affect pups’ behavior. Spectrograms of USVs were generated and analyzed by using Avisoft SASLab pro. Recording data were filtered in a range of 30-90 kHz, and start/end time, duration, peak frequency, and peak amplitude for each of the calls were collected as described previously (Tran et al., 2021). With the data extracted, the number of calls, average duration, average frequencies, and average amplitudes were determined for each pup individually and then averaged for each condition.

### 2.6. Tissue sample preparation and immunoblots

The cortex, hippocampus, and remaining portions of the forebrain (forebrain) were isolated from P10 each pup prenatally exposed to VPA for whole cell lysate analysis. Additionally, we used whole brain tissues from VPA-treated pups and isolated the postsynaptic density (PSD) fraction as shown previously (Farooq et al., 2016; Kim et al., 2016; Kim et al., 2018; Kim, Titcombe, et al., 2015; Kim, Violette, et al., 2015; Mendez-Vazquez et al., 2023; Shou et al., 2018; Zaytseva et al., 2023). Brain tissues were rinsed with PBS, collected to 15 ml tubes, homogenized by douncing in solution A, and spun at 2000 rpm for 10 min at 4°C. The supernatant (*whole cell lysates*) was centrifuged at 13000 rpm for 20 min at 4°C. The pellet (*P2*) was homogenized in solution B (320 mM Sucrose, 1 mM NaHCO_3_), placed on top of a 1 M sucrose and 1.2 M sucrose gradient, and centrifuged at 28000 rpm for 2 hours at 4°C. The resulting interface between the gradient was centrifuged at 40000 rpm for 45 min at 4°C after a six-fold dilution with solution B. The pellet (*synaptosome*) was resuspended in 1% Triton-X and 25 mM Tris buffer, inculcated for 30 min at 4°C while rocking, and spun at 13000 rpm for 30 min at 4°C. The pellet (*PSD*) was resuspended in 2% SDS and 25 mM Tris buffer. The protein concentration in each faction was determined by a BCA protein assay kit (Thermo Fisher Scientific, PI23227). Immunoblots were performed as described previously (Farooq et al., 2016; Kim et al., 2005; Kim et al., 2016; Kim, Sato, et al., 2015; Kim et al., 2018; Kim, Titcombe, et al., 2015; Kim, Violette, et al., 2015; Kim & Ziff, 2014; Roberts et al., 2021; Sathler et al., 2022; Sathler et al., 2021; Shou et al., 2018; Sun et al., 2019; Sztukowski et al., 2018; Tran et al., 2021). Equal amounts of protein samples were loaded on 10% glycine-SDS-PAGE gel. The gels were transferred to nitrocellulose membranes. The membranes were blocked (5% powdered milk) for 1 hour at room temperature, followed by overnight incubation with the primary antibodies at 4°C. Equal amounts of protein (whole cell lysates and PSD) were loaded on 10% SDS-PAGE gel and transferred to nitrocellulose membranes. Membranes were blotted with antibodies. The primary antibodies consisted of anti-δ-catenin (BD Biosciences, 1:1000, 611537), anti-GluA1 (RH95) (Millipore, 1:2000, MAB2263), anti-GluA2 (EPR18115) (Abcam, 1:2000, ab206293), anti-PSD-95 (K28/43) (NeuroMab, 1:1000, 75-028), and anti-actin (Abcam, 1:2000, ab3280) antibodies and developed with chemiluminescence (Life Tech, PI 34580). Protein bands were quantified using the NIH ImageJ software (https://imagej.nih.gov/ij/).

### 2.7. Immunohistochemistry

Immunohistochemistry was carried out as described previously (Tran et al., 2021). P5 VPA and saline-treated pups (both males and females) were subjected to 90 mins isolation from the nest. Pups in the nest were used as controls. Brains were then extracted and fixed with 4% paraformaldehyde for 3 days, and then 40 μm coronal sections were collected using a vibratome. The expression of the activity-regulated gene, c-Fos, was used to determine activation of neurons in the arcuate nucleus in the hypothalamus before or after isolation from the nest. Sections containing the arcuate nucleus in the hypothalamus were collected, permeabilized with diluting buffer (1% BSA, 0.4% Triton-X, 1% normal goat serum in pH 7.6 TBS), and blocked with 3% normal goat serum in TBS. The brain tissues were then incubated with anti-c-Fos antibody (Synaptic Systems, 226 003) diluted to 1:500 in a diluting buffer at 4°C for 18 hours. Afterwards, the sections were incubated for 2 hours with a secondary goat anti-rabbit IgG antibody conjugated with Alexa Fluor 647 (Invitrogen, A-21244). Nuclei were labeled by 4-,6-diamidino-2-phenylindole (DAPI). The sections were then mounted on microscope slides and coverslipped. Sections were imaged using the Olympus inverted microscope IX73. The Olympus cellSENS Dimensions software was used to measure c-Fos positive immunoreactivity on DAPI-positive cells in the hypothalamic region by colocalization analysis. Numbers of c-Fos-positive cells were then counted using particle analysis in the NIH ImageJ software (https://imagej.nih.gov/ij/).

### 2.8. Statistics

We used the GraphPad Prism 10 software to determine statistical significance (set at *p* < 0.05). Grouped results of single comparisons were tested for normality with the Shapiro-Wilk normality or Kolmogorov-Smirnov test and analyzed using the unpaired two-tailed Student’s t-test when data are normally distributed. Differences between multiple groups were assessed by two-way analysis of variance (ANOVA) with the Tukey test. The graphs were presented as mean ± Standard Deviation (SD).

## 3. Results

### 3.1. VPA prenatal exposure significantly decreases δ-catenin levels in the brains

In cultured human neurons, VPA treatment significantly decreases δ-catenin mRNA levels (Chanda et al., 2019). We thus examined whether VPA prenatal exposure significantly decreased δ-catenin protein levels in the brain of newborn mice. We intraperitoneally injected 250 mg/kg VPA or saline as a control (CTRL) to E12.5 pregnant mice. Following VPA injection, the brain tissues were collected from P10 pups. We then measured δ-catenin levels in whole cell lysates from the cortex, hippocampus, and forebrain using immunoblotting. When normalized to CTRL, VPA exposure significantly reduced δ-catenin levels in the cortex (CTRL, 1.000 ± 0.228 and VPA, 0.716 ± 0.273, *p* = 0.0129, t = 2.716, df = 21) (**Fig. 1a**), hippocampus (CTRL, 1.000 ± 0.456 and VPA, 0.583 ± 0.283, *p* = 0.0065, t = 2.957, df = 26) (**Fig. 1b**), and forebrain (CTRL, 1.000 ± 0.473 and VPA, 0.523 ± 0.256, *p* = 0.0011, t = 3.632, df = 28) (**Fig. 1c**). This demonstrates that prenatal exposure to VPA significantly decreases δ-catenin levels in the pups’ forebrains.

**Figure 1.**
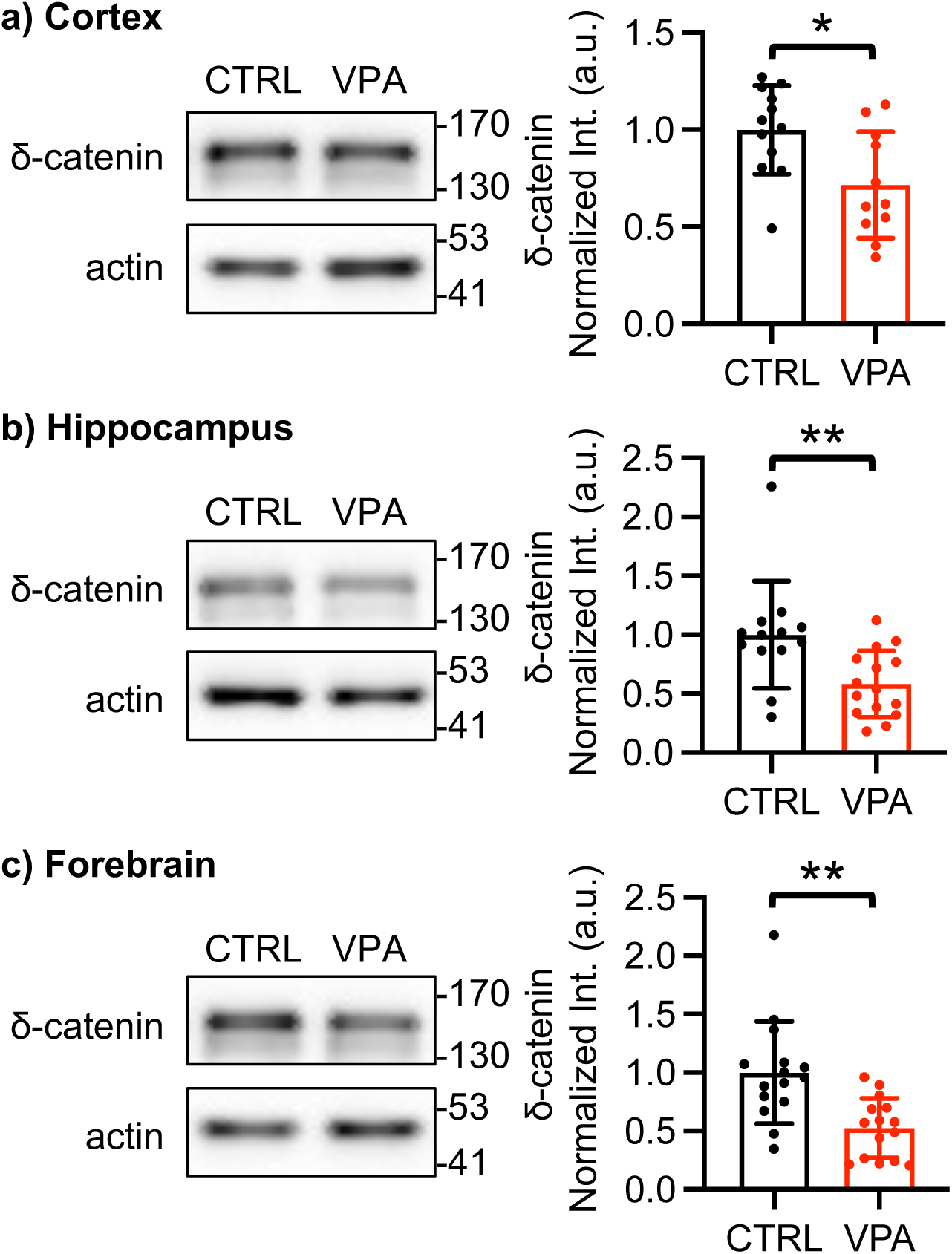
Representative immunoblots and summary graphs of normalized δ-catenin levels in **a)** the cortex (n = 12 CTRL and 11 VPA blots from 3 P10 pups), **b)** the hippocampus (n = 13 CTRL and 15 VPA blots from 3 P10 pups), and **c)** the forebrain (n = 15 CTRL and 15 VPA blots from 3 P10 pups. \**p* < 0.05 and \*\**p* < 0.01, the unpaired two-tailed Student’s t-test). The position of molecular mass markers (kDa) is shown on the right of the blots.

### 3.2. VPA treatment disrupts the development of dendritic branching, which is reversed by elevated **δ**-catenin expression

VPA exposure induces chronic impairments in dendritic morphology and AMPAR functions in developing human and mouse neurons (Chanda et al., 2019). The regulation of intracellular pathways by external signaling stimuli essentially controls dendrite development (Dong et al., 2015). In particular, the formation of dendritic morphology has been significantly linked to glutamate (Portera-Cailliau et al., 2003). This suggests that VPA-induced deficits in the dendritic development can be mediated by dysfunction in glutamatergic synapses. Notably, δ-catenin is important for synaptic structures and AMPAR function (Donta et al., 2022; Yuan & Arikkath, 2017). In fact, δ-catenin deficiency decreases dendritic arbor size, segment number, tip number, and branching complexity (Baumert et al., 2020; Israely et al., 2004; Matter et al., 2009). The acute loss of δ-catenin *in vitro* also impairs activity-dependent formation of spines (Brigidi et al., 2014). Furthermore, recent studies, including our own, suggest that δ-catenin deficiency induced by δ-catenin KO, an ASD-associated δ-catenin missense mutation, or δ-catenin knockdown significantly alters cortical neurons’ synaptic activity (Assendorp et al., 2022; Mendez-Vazquez et al., 2023). As prenatal exposure to VPA significantly reduces brain δ-catenin levels (**Fig. 1**), we hypothesized that the VPA effects on dendritic morphology were mediated by the loss of δ-catenin functions. To test this idea, we treated 4 DIV cultured mouse cortical neurons with 1 mM VPA for 3 days and analyzed the complexity of dendritic branching in developing neurons at 7 DIV using Sholl analysis (**Fig. 2**). In developing neurons treated with VPA, dendritic branching was significantly reduced when compared with control neurons (CTRL) (**Fig. 2** **and Table 1**). When of δ-catenin was exogenously expressed, dendritic branching was significantly increased in the proximal area (< 60 μm from soma) compared with CTRL (**Fig. 2** **and Table 1**), which is consistent with the previous findings (Arikkath et al., 2008; Baumert et al., 2020; Martinez et al., 2003). This result suggests that dendrite branching is modulated by δ-catenin. Importantly, we found that reduced dendritic branching was markedly reversed by δ-catenin expression in the distal area (> 80 μm from soma) of VPA-treated neurons (**Fig. 2** **and Table 1**). This suggests that the loss of δ-catenin functions contributes to VPA-induced deficits in neuronal development.

**Figure 2.**
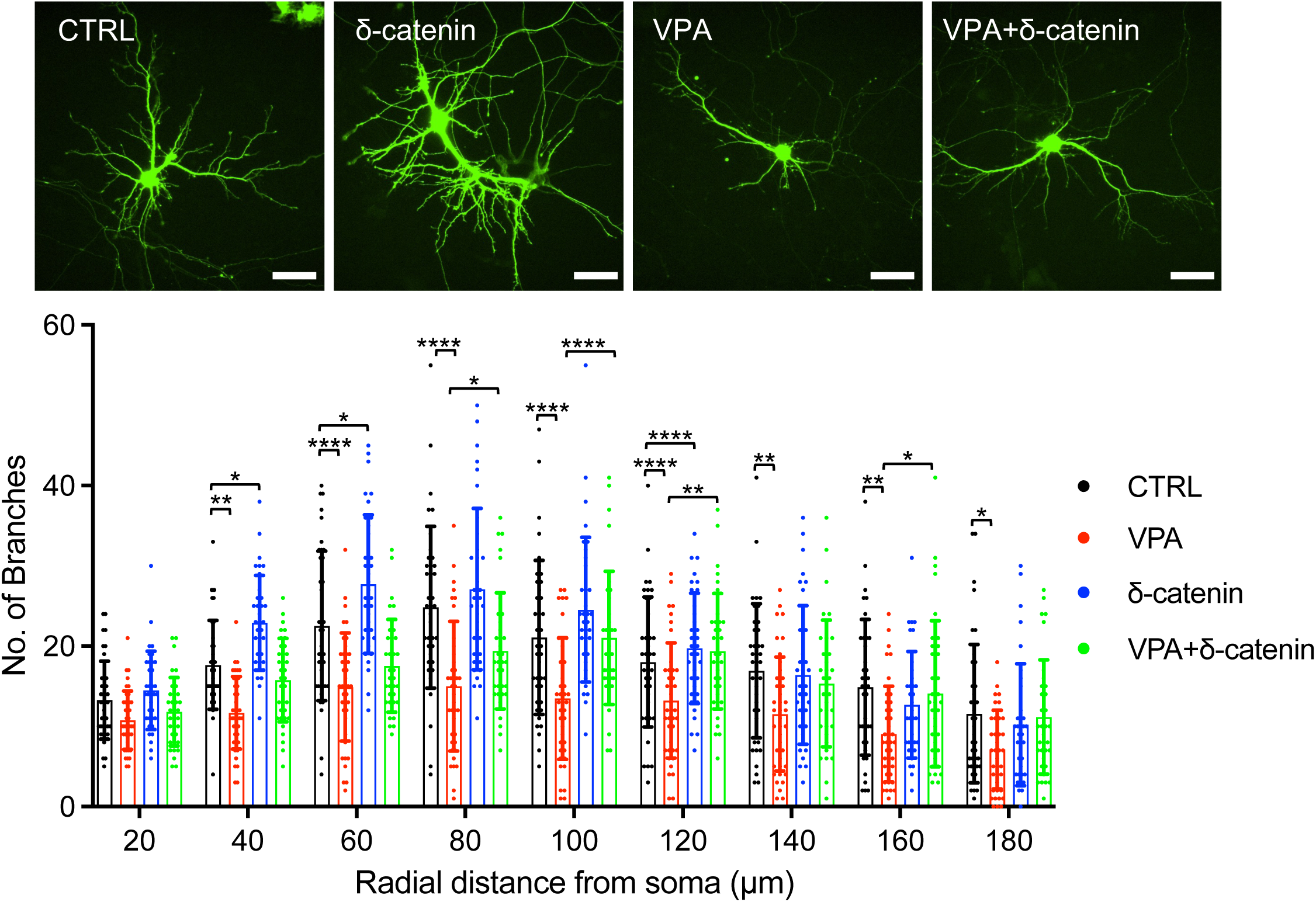
Representative images of dendritic branches in cultured cortical neurons in each condition. Sholl analysis of dendrite morphology for each condition (n = number of neurons. CTRL = 39, δ-catenin = 40, VPA = 35, and VPA + δ-catenin = 35. For VPA vs VPA + δ-catenin. \**p* < 0.05, \*\**p* < 0.01, and \*\*\*\**p* < 0.0001, Two-way ANOVA with the Tukey test). A scale bar indicates 50 μm.

**Table 1.** The summary of Sholl analysis.

| Radial distance from soma ($\mu\text{m}$) | Number of branches | | | |
| --- | --- | --- | --- | --- |
| | CTRL | VPA | δ-catenin | VPA + δ-catenin |
| 20 | 13.282 ± 4.856 | 10.775 ± 3.634 | 14.486 ± 4.889 | 11.829 ± 4.253 |
| | CTRL vs. VPA, $p = 0.4320$, CTRL vs. δ-catenin, $p = 0.8969$, and VPA vs. VPA + δ-catenin, $p = 0.9268$ | | | |
| 40 | 17.667 ± 5.527 | 11.700 ± 4.513 | 22.914 ± 5.888 | 15.771 ± 5.197 |
| | CTRL vs. VPA, $p = 0.0019$, CTRL vs. δ-catenin, $p = 0.0123$, and VPA vs. VPA + δ-catenin, $p = 0.0808$ | | | |
| 60 | 22.513 ± 9.319 | 14.900 ± 6.732 | 27.714 ± 8.635 | 17.543 ± 5.762 |
| | CTRL vs. VPA, $p < 0.0001$, CTRL vs. δ-catenin, $p = 0.0134$, and VPA vs. VPA + δ-catenin, $p = 0.4096$ | | | |
| 80 | 24.846 ± 10.077 | 15.000 ± 8.077 | 27.086 ± 10.057 | 19.400 ± 7.244 |
| | CTRL vs. VPA, $p < 0.0001$, CTRL vs. δ-catenin, $p = 0.5610$, and VPA vs. VPA + δ-catenin, $p = 0.0496$ | | | |
| 100 | 21.077 ± 9.590 | 13.475 ± 7.572 | 24.543 ± 9.014 | 21.029 ± 8.301 |
| | CTRL vs. VPA, $p < 0.0001$, CTRL vs. δ-catenin, $p = 0.1822$, and VPA vs. VPA + δ-catenin, $p < 0.0001$ | | | |
| 120 | 18.026 ± 8.100 | 13.225 ± 7.174 | 19.743 ± 6.879 | 19.371 ± 7.203 |
| | CTRL vs. VPA, $p < 0.0001$, CTRL vs. δ-catenin, $p < 0.0001$, and VPA vs. VPA + δ-catenin, $p = 0.0019$ | | | |
| 140 | 16.923 ± 8.368 | 11.550 ± 7.107 | 16.400 ± 8.630 | 15.343 ± 7.904 |
| | CTRL vs. VPA, $p = 0.0068$, CTRL vs. δ-catenin, $p = 0.9902$, and VPA vs. VPA + δ-catenin, $p = 0.1182$ | | | |
| 160 | 14.872 ± 8.449 | 9.050 ± 5.944 | 12.686 ± 6.632 | 14.086 ± 9.099 |
| | CTRL vs. VPA, $p = 0.0027$, CTRL vs. δ-catenin, $p = 0.5809$, and VPA vs. VPA + δ-catenin, $p = 0.0171$ | | | |
| 180 | 11.590 ± 8.620 | 7.125 ± 4.889 | 10.229 ± 7.593 | 11.171 ± 7.123 |
| | CTRL vs. VPA, $p = 0.0365$, CTRL vs. δ-catenin, $p = 0.8581$, and VPA vs. VPA + δ-catenin, $p = 0.0837$ | | | |

Two-way ANOVA with the Tukey test
| ANOVA table | SS (Type III) | DF | MS | F (DFn, DFd) | $p$ value |
| --- | --- | --- | --- | --- | --- |
| Interaction | 3891 | 24 | 162.1 | F (24, 1305) = 2.976 | $p < 0.0001$ |
| Row Factor | 19420 | 8 | 2428 | F (8, 1305) = 44.57 | $p < 0.0001$ |
| Column Factor | 11231 | 3 | 3744 | F (3, 1305) = 68.73 | $p < 0.0001$ |
| Residual | 71086 | 1305 | 54.47 |  |  |

### 3.3. VPA prenatal exposure significantly reduces synaptic AMPAR levels in pups

As we confirmed that VPA significantly reduced δ-catenin levels in pups’ brains (**Fig. 1**), we wanted to understand how VPA affected synaptic AMPAR levels in these animals. The PSD fractions of entire forebrains from saline (CTRL) or prenatally VPA-exposed P10 pups (males and females) were collected and analyzed using immunoblotting. We found a significant reduction in synaptic δ-catenin (CTRL, 1.000 ± 0.241 and VPA, 0.105 ± 0.071, *p* < 0.0001, t = 10.67, df = 16.00), GluA1 (CTRL, 1.000 ± 0.397 and VPA, 0.538 ± 0.243, *p* < 0.0089, t = 2.983, df = 16.00), GluA2 (CTRL, 1.000 ± 0.074 and VPA, 0.7025 ± 0.290, *p* < 0.0086, t = 2.991, df = 16.00), and PSD95 (CTRL, 1.000 ± 0.425 and VPA, 0.217 ± 0.125, *p* < 0.0001, t = 5.302, df = 16.00) (**Fig. 3**). This demonstrates that VPA prenatal exposure markedly decreases synaptic AMPAR levels and structural proteins in newly born mice, which is likely mediated by the loss of δ-catenin functions. This further suggests that pups prenatally exposed to VPA have altered glutamatergic synaptic transmission.

**Figure 3.**
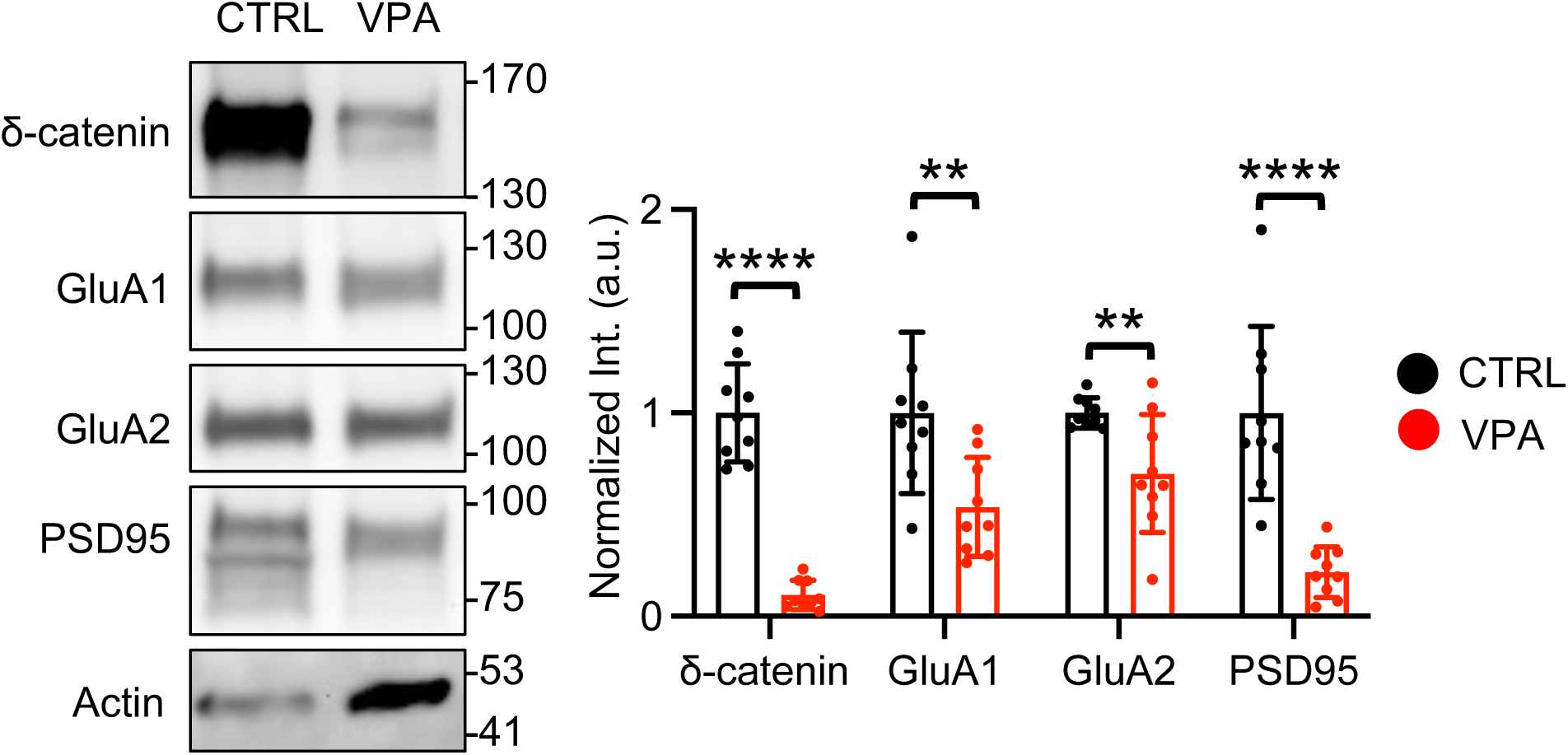
Representative immunoblots and summary graphs of normalized synaptic proteins levels in entire forebrains of P10 pups in each condition (n = 9 immunoblots from 3 animals in each condition, \*\**p* < 0.01 and \*\*\*\**p* < 0.0001, the unpaired two-tailed Student’s t-test). The position of molecular mass markers (kDa) is shown on the right of the blots.

### 3.4. Prenatal exposure to VPA impairs ultrasonic vocalization in newly born pups

Ultrasonic vocalizations (USVs) are high frequency vocalizations used by rodents in male-female, juvenile, and mother-infant interactions (Branchi et al., 2001; Fischer & Hammerschmidt, 2011; Hahn & Lavooy, 2005; Portfors, 2007). When a pup experiences an unpleasant stimulus, like being separated from their nest, they will vocalize between 30 and 90 kHz, which will cause the mother to respond by retrieving them, an indication of animals’ communication (Ferhat et al., 2016; Hofer, 1996; Noirot, 1966; Scattoni et al., 2009; Sewell, 1970; Zimprich et al., 2017). We thus measured USVs to address whether prenatal exposure to VPA affected animals’ communication, a key pathology in ASD. 90 min after isolation from the nest, we recorded USVs from P5-7 pups of control (CTRL) and VPA-exposed mice. As mouse pup USVs occur in the 40-80 kHz range (Ehret & Haack, 1981), spectrograms in this range for each recording were created and analyzed as shown previously (Tran et al., 2021). We found that pups prenatally exposed to VPA emitted less vocalizations than control animals during the 5 min testing period (CTRL, 250.2 ± 137.1 and VPA, 142.5 ± 115.6, *p* = 0.0035, t = 3.063, df = 50) (**Fig. 4a**). The total duration of calls of VPA-exposed pups was also significantly reduced compared with WT and HET mice (CTRL, 8.888 ± 15.424 sec and VPA, 5.417 ± 4.479 sec, *p* = 0.0151, t = 2.516, df = 50) (**Fig. 4b**). We also analyzed the peak frequency of USVs and found that VPA-exposed pups produced higher frequency USVs than CTRL (CTRL, 63.96 ± 2.232 KHz and VPA, 65.78 ± 2.358 KHz, *p* = 0.0073, t = 2.798, df = 50) (**Fig. 4c**). When comparing the average duration of each USV call from each condition, there was no significant difference (CTRL, 34.15 ± 25.179 ms and VPA, 37.22 ± 9.266 KHz, *p* = 0.1470, t = 1.473, df = 50) (**Fig. 4d**). Finally, we found no difference in the peak amplitude of calls in each group (CTRL, -20.31 ± 2.004 dB and VPA, -19.89 ± 3.937 dB, *p* = 0.16247, t = 0.4923, df = 50) (**Fig. 4e**). This demonstrates that VPA-exposed pups generated less but high frequency calls, which suggests that prenatal exposure of VPA disrupts animals’ communication, which is consistent with the previous findings (Felix-Ortiz & Febo, 2012; Gzielo et al., 2020; Potasiewicz et al., 2020; Tsuji et al., 2020).

**Figure 4.**
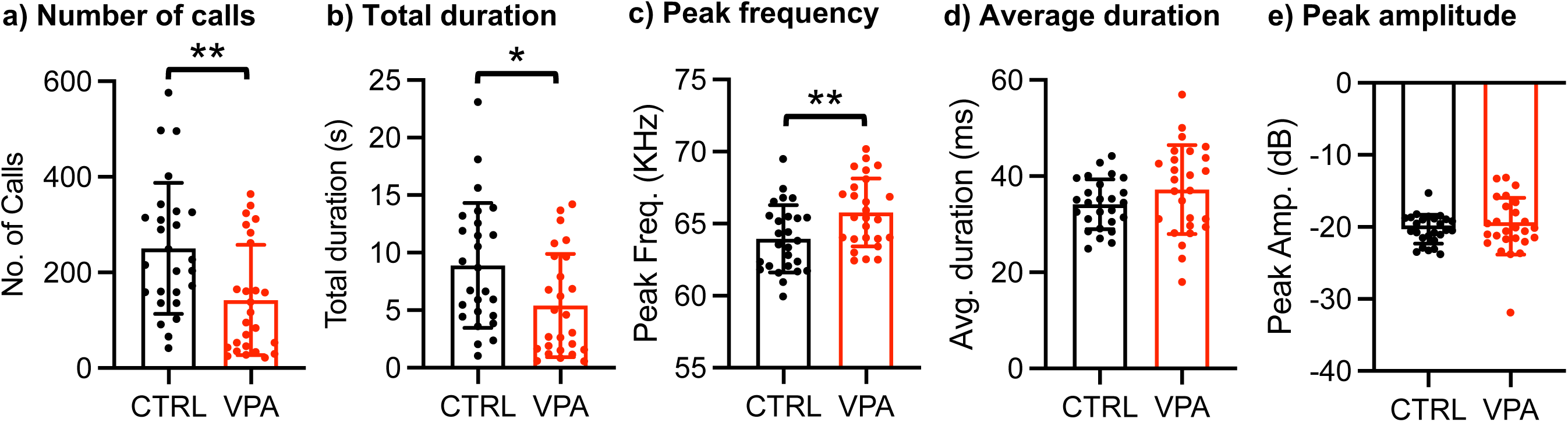
Abnormal ultrasonic vocalizations in isolated pups prenatally exposed to VPA relative to control pups. **a)** Total number of USVs emitted, **b)** duration of individual calls, **c)** peak frequency, **d)** peak amplitude (or intensity), and **e)** total duration of calls in each genotype during 5-minute isolation period. (n = number of animals, CTRL = 26 and VPA =26 mice, \**p* < 0.05 and \*\**p* < 0.01, the unpaired two-tailed Student’s t-test). Mean ± SD.

### 3.5. Reduction of neuronal activation in the arcuate nucleus of VPA-exposed animals following isolation

When pups are separated from the nest, their body temperatures drop, which in turn causes USV emissions (Noirot, 1966). A study reveals that after being separated from the nest for 90 minutes, the pups’ body temperature changes cause a substantial activation of agouti-related peptide (AgRP) neurons in the arcuate nucleus of the hypothalamus, which is shown to induce USVs (Zimmer et al., 2019). Therefore, after being separated from their nest, pups’ activation of AgRP neurons in the arcuate nucleus is regulated by thermal insulation and is linked to the emission of USVs (Zimmer et al., 2019). δ-catenin is known to regulate neuronal activity by controlling synaptic AMPAR expression (Assendorp et al., 2022; Farooq et al., 2016; Kosik et al., 2005; Mendez-Vazquez et al., 2023; Restituito et al., 2011; Silverman et al., 2007; Yuan & Arikkath, 2017; Yuan et al., 2015). Additionally, we reveal that prenatal exposure of VPA significantly reduce synaptic δ-catenin and AMPAR levels (**Fig. 3**). Therefore, we hypothesized that VPA can alter neuronal activation in the arcuate nucleus following isolation from the nest, which in turn affects USVs. We thus measured expression of the activity-regulated gene, c-Fos, as a neuronal activity marker (Bullitt, 1990; Greenberg & Ziff, 1984) to determine activation of arcuate nucleus neurons before or after isolation (**Fig. 5**). To quantify c-Fos immunoreactivity at the level of individual neurons, nuclei were labeled with DAPI, and we measured c-Fos signals on individual DAPI-positive cells in the arcuate nucleus. Saline-exposed mice after a 90 min isolation exhibited significantly increased neuronal activity in the hypothalamus compared to mice in the nest (**Fig. 5** **and Table 2**), which is consistent with the previous reports (Tran et al., 2021; Zimmer et al., 2019). Interestingly, basal c-Fos immunoreactivity in the arcuate nucleus of VPA-exposed pups in the nest was significantly higher compared to saline-exposed animals (**Fig. 5** **and Table 2**), which is consistent with the previous findings (Campolongo et al., 2018; Dubiel & Kulesza, 2016; Zappala et al., 2023). However, a 90 min isolation was unable to further elevate c-Fos immunoreactivity in the arcuate nucleus of VPA-exposed mice compared to nest control animals (**Fig. 5** **and Table 2**). These results suggest that prenatal exposure of VPA is unable to increase neuronal activity in the arcuate nucleus following isolation, which likely underlies altered USVs in these animals.

**Figure 5.**
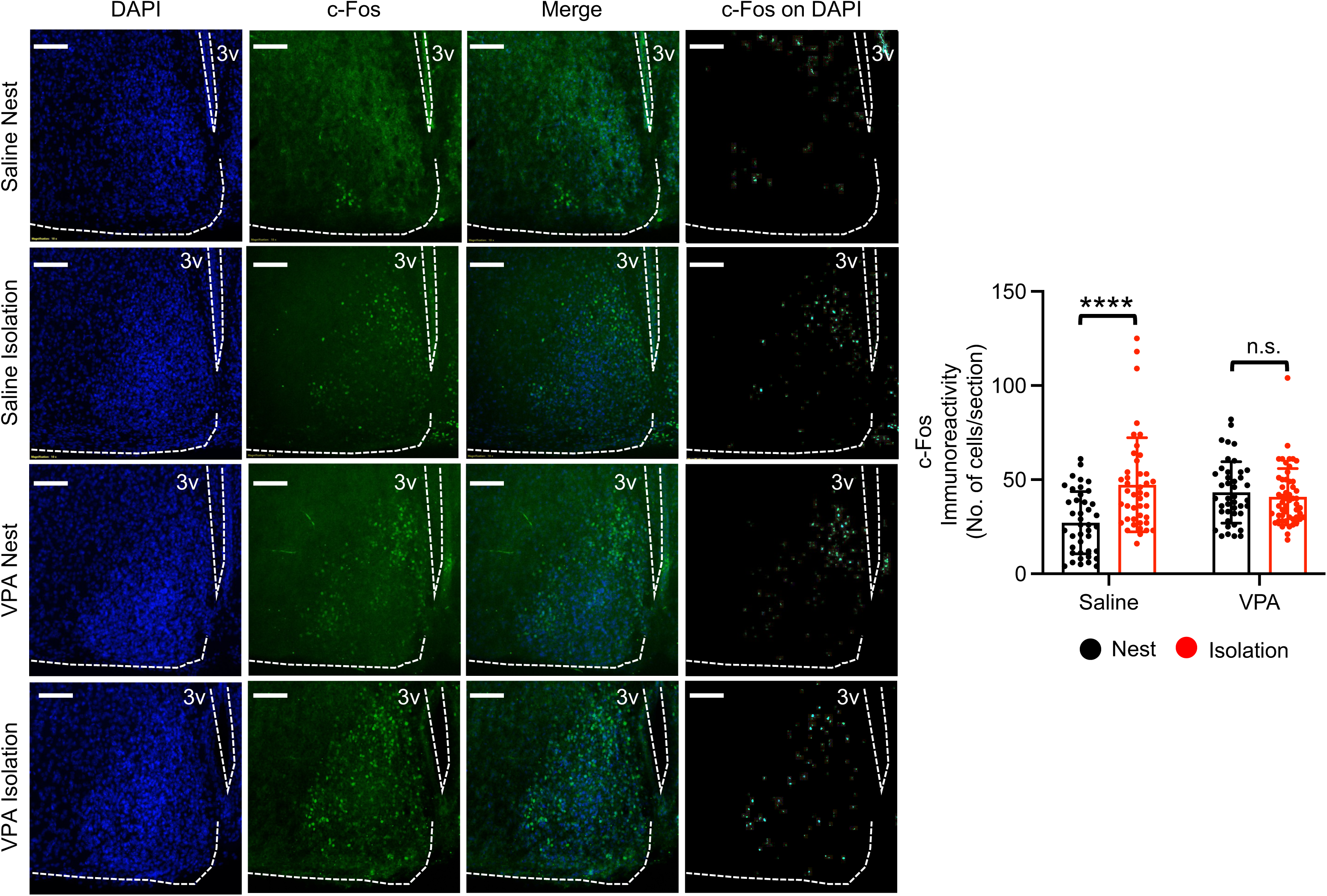
Left panels: Representative c-Fos immunoreactivity (Green), DAPI (Blue), merge images, and c-Fos signals colocalized with DAPI images in each condition. The graph on the right shows that neuronal activation is significantly reduced in the VPA-treated arcuate nucleus following isolation from the nest (n = Saline Nest; 41 sections from 4 animals, Saline Isolation; 43 sections from 4 animals, VPA Nest; 42 sections from 5 animal, and VPA Isolation; 53 sections from 5 animals, \*\*\*\**p* < 0.0001, Two-way ANOVA with the Tukey test). A scale bar indicates 100 μm. 3v, the third ventricle. n.s. indicates not significant.

**Table 2.**
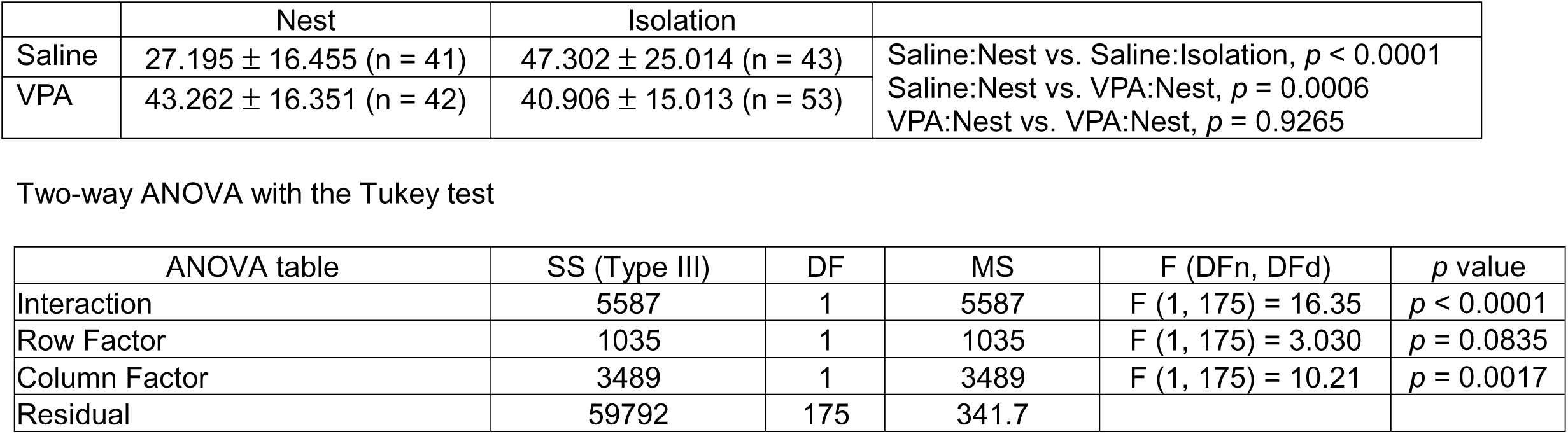
The summary of c-Fos immunoreactivity analysis.

## 4. Discussion

VPA is a medication primarily used to treat epilepsy and bipolar disorder. It has been suggested to be an anticonvulsant and mood-stabilizing drug that works by affecting certain neurotransmitters in the brain (Rahman et al., 2023). Some research has explored the potential link between VPA exposure during pregnancy and an increased risk of ASD in the offspring. However, it’s important to note that this association is complex and not fully understood. In the current study, we focus on the role of δ-catenin to link prenatal exposure to VPA and ASD because: 1) VPA treatment significantly reduces δ-catenin mRNA levels, 2) a loss of δ-catenin functions phenocopies the VPA effects on neuronal development, and 3) δ-catenin deficiency in humans causes severe ASD. The current study has discovered that: 1) prenatal exposure of VPA in mice significantly decreases total δ-catenin protein levels in the pups’ brains (**Fig. 1**), 2) VPA treatment markedly disrupts dendritic branching in cultured cortical neurons, which is reversed by increased δ-catenin expression (**Fig. 2**), 3) prenatal exposure to VPA significantly decreases synaptic δ-catenin and AMPAR expression in newly born pups (**Fig. 3**), 4) VPA-exposed pups exhibit impaired USVs (**Fig. 4**), and 5) isolation from the nest is unable to activate neurons in the arcuate nucleus, which is required for USVs, following prenatal exposure to VPA (**Fig. 5**). Our results, together with existing data, indicate that prenatal exposure to VPA significantly lowers synaptic δ-catenin levels, disrupting dendritic development and AMPAR-mediated glutamatergic synaptic activity in neurons, including cells in the arcuate nucleus. This ultimately disrupts animals’ communication (e.g. USVs), a key pathology in ASD. Hence, our work suggests that the loss of δ-catenin functions underlies VPA-induced ASD in early childhood.

The δ*-catenin* gene is categorized as a ‘strong candidate’ regarding its association with ASD in the Simons Foundation Autism Research Initiative (SFARI) Database. δ-catenin expression is closely linked to other ASD-risk genes involved in synaptic structure and function, such as GluA2, cadherins, GRIP, and synaptic Ras GTPase activating protein 1 (SYNGAP1), further implying its potential roles in ASD etiology (Assendorp et al., 2022; El-Amraoui & Petit, 2010; Mejias et al., 2011; Ramanathan et al., 2004). Furthermore, the Cri-du-chat syndrome results from variable hemizygous deletions in the short arm of chromosome 5p, including the δ*-catenin* gene. The clinical features comprise growth delay, severe mental retardation, and speech delay (Niebuhr, 1978; Overhauser et al., 1986). In those individuals with the syndrome who have mild or no obvious intellectual disability, the deletion often does not include the δ*-catenin* gene (Medina et al., 2000). Additionally, individuals carrying a partial deletion and a partial duplication of the δ*-catenin* gene exhibit the Cri-du-chat syndrome, but only mild cognitive disability (Sardina et al., 2014). This suggests the δ*-catenin* gene plays crucial roles in etiology of the syndrome. Approximately 40% of individuals with this syndrome show some autistic-like characteristics (Moss et al., 2008). This is considerably higher than expected in the general population (Moss et al., 2008). Importantly, genetic studies identify loss of δ-catenin function as a significant factor for both the Cri-du-chat syndrome and ASD (Harvard et al., 2005; Miller et al., 2019; Weiss et al., 2009; Zhang et al., 2016). δ-catenin mutations also have been implicated in many other neurological disorders, including cerebral palsy, schizophrenia, anxiety disorders and Alzheimer’s disease (Jun et al., 2012; McMichael et al., 2014; Nivard et al., 2014; Vrijenhoek et al., 2008). Therefore, comprehending the loss of δ-catenin functions can lead to an understanding of shared etiology and pathophysiology among these diseases.

There may be many different factors that make a child more likely to have ASD, including environmental and genetic factors (Christensen et al., 2013; Huguet et al., 2013). The interaction of both environmental and genetic risk factors for ASD is an important area of investigation as the complementary contribution of the two factors may enhance the etiologic strength in ASD pathogenesis (Mabunga et al., 2015). However, the molecular link of environmental and genetic risk factors for ASD has not been fully understood (Chaste & Leboyer, 2012). Our findings suggest that the loss of δ-catenin functions is likely the common ASD pathological target arising from both genetic and environmental factors, which is one of the important aspects of the current research.

## 5. Conclusions

Pregnant women who had VPA treatment have a higher prevalence of ASD in their children. However, our current knowledge of the mechanism of VPA-induced ASD is far from complete. VPA treatment significantly reduces δ-catenin mRNA levels in cultured human neurons. Moreover, a loss of δ-catenin functions phenocopies VPA-induced disruptions in glutamatergic synapses. Here, we discover that VPA exposure can impair glutamatergic synapses via a reduction of δ-catenin expression, results in altered neuronal activity in the hypothalamus and ultimately deficits in animals’ communication, a key pathological change in ASD. This thus suggests that a loss of δ-catenin functions is likely to be the molecular mechanism underlying VPA-induced ASD.

## Supporting information

Untruncated image of gels

## Acknowledgements

We thank members of the Kim laboratory for their generous support. We appreciate thoughtful suggestions from Dr. Soham Chanda. This work is funded by College Research Council Shared Research Program, Student Experiential Learning Grants from Colorado State University, the Jerome Lejeune Foundation, and the NIH grant (1R01MH132921) for SK.

## References

Al Sagheer, T., Haida, O., Balbous, A., Francheteau, M., Matas, E., Fernagut, P. O., & Jaber, M. (2018). Motor Impairments Correlate with Social Deficits and Restricted Neuronal Loss in an Environmental Model of Autism. Int J Neuropsychopharmacol, 21(9), 871–882. 10.1093/ijnp/pyy043

Arikkath, J., Israely, I., Tao, Y., Mei, L., Liu, X., & Reichardt, L. F. (2008). Erbin controls dendritic morphogenesis by regulating localization of delta-catenin. The Journal of neuroscience : the official journal of the Society for Neuroscience, 28(28), 7047–7056. 10.1523/JNEUROSCI.0451-08.2008

Assendorp, N., Depp, M., Fossati, M., Dingli, F., Loew, D., & Charrier, C. (2022). CTNND2 moderates neuronal excitation and links human evolution to prolonged synaptic development in the neocortex. bioRxiv, 2022.2009.2013.507776. 10.1101/2022.09.13.507776

Baumert, R., Ji, H., Paulucci-Holthauzen, A., Wolfe, A., Sagum, C., Hodgson, L., … McCrea, P. D. (2020). Novel phospho-switch function of delta-catenin in dendrite development. J Cell Biol, 219(11). 10.1083/jcb.201909166

Betancur, C., Sakurai, T., & Buxbaum, J. D. (2009). The emerging role of synaptic cell-adhesion pathways in the pathogenesis of autism spectrum disorders. Trends Neurosci, 32(7), 402–412. 10.1016/j.tins.2009.04.003

Bourgeron, T. (2015). From the genetic architecture to synaptic plasticity in autism spectrum disorder. Nature reviews. Neuroscience, 16(9), 551–563. 10.1038/nrn3992

Branchi, I., Santucci, D., & Alleva, E. (2001). Ultrasonic vocalisation emitted by infant rodents: a tool for assessment of neurobehavioural development. Behav Brain Res, 125(1-2), 49–56. 10.1016/s0166-4328(01)00277-7

Brigidi, G. S., Sun, Y., Beccano-Kelly, D., Pitman, K., Mobasser, M., Borgland, S. L., … Bamji, S. X. (2014). Palmitoylation of delta-catenin by DHHC5 mediates activity-induced synapse plasticity. Nature neuroscience, 17(4), 522–532. 10.1038/nn.3657

Brown, N. A., Kao, J., & Fabro, S. (1980). Teratogenic potential of valproic acid. Lancet, 1(8169), 660–661. https://www.ncbi.nlm.nih.gov/pubmed/6102670

Bullitt, E. (1990). Expression of c-fos-like protein as a marker for neuronal activity following noxious stimulation in the rat. J Comp Neurol, 296(4), 517–530. 10.1002/cne.902960402

Campolongo, M., Kazlauskas, N., Falasco, G., Urrutia, L., Salgueiro, N., Hocht, C., & Depino, A. M. (2018). Sociability deficits after prenatal exposure to valproic acid are rescued by early social enrichment. Mol Autism, 9, 36. 10.1186/s13229-018-0221-9

Chaliha, D., Albrecht, M., Vaccarezza, M., Takechi, R., Lam, V., Al-Salami, H., & Mamo, J. (2020). A Systematic Review of the Valproic-Acid-Induced Rodent Model of Autism. Dev Neurosci, 42(1), 12–48. 10.1159/000509109

Chanda, S., Ang, C. E., Lee, Q. Y., Ghebrial, M., Haag, D., Shibuya, Y., … Sudhof, T. C. (2019). Direct Reprogramming of Human Neurons Identifies MARCKSL1 as a Pathogenic Mediator of Valproic Acid-Induced Teratogenicity. Cell Stem Cell. 10.1016/j.stem.2019.04.021

Chaste, P., & Leboyer, M. (2012). Autism risk factors: genes, environment, and gene-environment interactions. Dialogues Clin Neurosci, 14(3), 281–292. 10.31887/DCNS.2012.14.3/pchaste

Christensen, J., Gronborg, T. K., Sorensen, M. J., Schendel, D., Parner, E. T., Pedersen, L. H., & Vestergaard, M. (2013). Prenatal valproate exposure and risk of autism spectrum disorders and childhood autism. JAMA, 309(16), 1696–1703. 10.1001/jama.2013.2270

Chung, L., Bey, A. L., & Jiang, Y. H. (2012). Synaptic plasticity in mouse models of autism spectrum disorders. Korean J Physiol Pharmacol, 16(6), 369–378. 10.4196/kjpp.2012.16.6.369

Deykin, E. Y., & MacMahon, B. (1979). Viral exposure and autism. Am J Epidemiol, 109(6), 628–638. 10.1093/oxfordjournals.aje.a112726

Dong, X., Shen, K., & Bulow, H. E. (2015). Intrinsic and extrinsic mechanisms of dendritic morphogenesis. Annu Rev Physiol, 77, 271–300. 10.1146/annurev-physiol-021014-071746

Donta, M. S., Srivastava, Y., & McCrea, P. D. (2022). Delta-Catenin as a Modulator of Rho GTPases in Neurons. Front Cell Neurosci, 16, 939143. 10.3389/fncel.2022.939143

Dubiel, A., & Kulesza, R. J., Jr. (2016). Prenatal valproic acid exposure disrupts tonotopic c-Fos expression in the rat brainstem. Neuroscience, 324, 511–523. 10.1016/j.neuroscience.2016.01.030

Ehret, G., & Haack, B. (1981). Categorical perception of mouse pup ultrasound by lactating females. Naturwissenschaften, 68(4), 208–209. 10.1007/BF01047208

Eidelman, A. I., & Samueloff, A. (2002). The pathophysiology of the fetus of the diabetic mother. Semin Perinatol, 26(3), 232–236. 10.1053/sper.2002.34215

El-Amraoui, A., & Petit, C. (2010). Cadherins as targets for genetic diseases. Cold Spring Harbor perspectives in biology, 2(1), a003095. 10.1101/cshperspect.a003095

Farooq, M., Kim, S., Patel, S., Khatri, L., Hikima, T., Rice, M. E., & Ziff, E. B. (2016). Lithium increases synaptic GluA2 in hippocampal neurons by elevating the delta-catenin protein. Neuropharmacology. 10.1016/j.neuropharm.2016.10.025

Felix-Ortiz, A. C., & Febo, M. (2012). Gestational valproate alters BOLD activation in response to complex social and primary sensory stimuli. PLoS One, 7(5), e37313. 10.1371/journal.pone.0037313

Ferhat, A. T., Torquet, N., Le Sourd, A. M., de Chaumont, F., Olivo-Marin, J. C., Faure, P., … Ey, E. (2016). Recording Mouse Ultrasonic Vocalizations to Evaluate Social Communication. J Vis Exp(112). 10.3791/53871

Fischer, J., & Hammerschmidt, K. (2011). Ultrasonic vocalizations in mouse models for speech and socio-cognitive disorders: insights into the evolution of vocal communication. Genes Brain Behav, 10(1), 17–27. 10.1111/j.1601-183X.2010.00610.x

Gilbert, J., & Man, H. Y. (2016). The X-Linked Autism Protein KIAA2022/KIDLIA Regulates Neurite Outgrowth via N-Cadherin and delta-Catenin Signaling. eNeuro, 3(5). 10.1523/ENEURO.0238-16.2016

Greenberg, M. E., & Ziff, E. B. (1984). Stimulation of 3T3 cells induces transcription of the c-fos proto-oncogene. Nature, 311(5985), 433–438. 10.1038/311433a0

Guo, H., Wang, T., Wu, H., Long, M., Coe, B. P., Li, H., … Xia, K. (2018). Inherited and multiple de novo mutations in autism/developmental delay risk genes suggest a multifactorial model. Mol Autism, 9, 64. 10.1186/s13229-018-0247-z

Gzielo, K., Potasiewicz, A., Holuj, M., Litwa, E., Popik, P., & Nikiforuk, A. (2020). Valproic acid exposure impairs ultrasonic communication in infant, adolescent and adult rats. Eur Neuropsychopharmacol, 41, 52–62. 10.1016/j.euroneuro.2020.09.006

Hahn, M. E., & Lavooy, M. J. (2005). A review of the methods of studies on infant ultrasound production and maternal retrieval in small rodents. Behav Genet, 35(1), 31–52. 10.1007/s10519-004-0854-7

Hampson, D. R., Gholizadeh, S., & Pacey, L. K. (2012). Pathways to drug development for autism spectrum disorders. Clin Pharmacol Ther, 91(2), 189–200. 10.1038/clpt.2011.245

Harvard, C., Malenfant, P., Koochek, M., Creighton, S., Mickelson, E. C., Holden, J. J., … Rajcan-Separovic, E. (2005). A variant Cri du Chat phenotype and autism spectrum disorder in a subject with de novo cryptic microdeletions involving 5p15.2 and 3p24.3-25 detected using whole genomic array CGH. Clin Genet, 67(4), 341–351. 10.1111/j.1399-0004.2005.00406.x

Hofer, M. A. (1996). Multiple regulators of ultrasonic vocalization in the infant rat. Psychoneuroendocrinology, 21(2), 203–217. 10.1016/0306-4530(95)00042-9

Huguet, G., Ey, E., & Bourgeron, T. (2013). The genetic landscapes of autism spectrum disorders. Annu Rev Genomics Hum Genet, 14, 191–213. 10.1146/annurev-genom-091212-153431

Israely, I., Costa, R. M., Xie, C. W., Silva, A. J., Kosik, K. S., & Liu, X. (2004). Deletion of the neuron-specific protein delta-catenin leads to severe cognitive and synaptic dysfunction. Curr Biol, 14(18), 1657–1663. 10.1016/j.cub.2004.08.065

Ito, T., Ando, H., Suzuki, T., Ogura, T., Hotta, K., Imamura, Y., … Handa, H. (2010). Identification of a primary target of thalidomide teratogenicity. Science, 327(5971), 1345–1350. 10.1126/science.1177319

Jentink, J., Loane, M. A., Dolk, H., Barisic, I., Garne, E., Morris, J. K., … Group, E. A. S. W. (2010). Valproic acid monotherapy in pregnancy and major congenital malformations. N Engl J Med, 362(23), 2185–2193. 10.1056/NEJMoa0907328

Jun, G., Moncaster, J. A., Koutras, C., Seshadri, S., Buros, J., McKee, A. C., … Farrer, L. A. (2012). delta-Catenin is genetically and biologically associated with cortical cataract and future Alzheimer-related structural and functional brain changes. PLoS One, 7(9), e43728. 10.1371/journal.pone.0043728

Karimi, P., Kamali, E., Mousavi, S. M., & Karahmadi, M. (2017). Environmental factors influencing the risk of autism. J Res Med Sci, 22, 27. 10.4103/1735-1995.200272

Kataoka, S., Takuma, K., Hara, Y., Maeda, Y., Ago, Y., & Matsuda, T. (2013). Autism-like behaviours with transient histone hyperacetylation in mice treated prenatally with valproic acid. Int J Neuropsychopharmacol, 16(1), 91–103. 10.1017/S1461145711001714

Kim, S., Lapham, A. N., Freedman, C. G., Reed, T. L., & Schmidt, W. K. (2005). Yeast as a tractable genetic system for functional studies of the insulin-degrading enzyme. J Biol Chem, 280(30), 27481–27490. 10.1074/jbc.M414192200

Kim, S., Pick, J. E., Abera, S., Khatri, L., Ferreira, D. D., Sathler, M. F., … Ziff, E. B. (2016). Brain region-specific effects of cGMP-dependent kinase II knockout on AMPA receptor trafficking and animal behavior. Learn Mem, 23(8), 435–441. 10.1101/lm.042960.116

Kim, S., Sato, Y., Mohan, P. S., Peterhoff, C., Pensalfini, A., Rigoglioso, A., … Nixon, R. A. (2015). Evidence that the rab5 effector APPL1 mediates APP-betaCTF-induced dysfunction of endosomes in Down syndrome and Alzheimer’s disease. Mol Psychiatry. 10.1038/mp.2015.97

Kim, S., Shou, J., Abera, S., & Ziff, E. B. (2018). Sucrose withdrawal induces depression and anxiety-like behavior by Kir2.1 upregulation in the nucleus accumbens. Neuropharmacology, 130, 10–17. 10.1016/j.neuropharm.2017.11.041

Kim, S., Titcombe, R. F., Zhang, H., Khatri, L., Girma, H. K., Hofmann, F., … Ziff, E. B. (2015). Network compensation of cyclic GMP-dependent protein kinase II knockout in the hippocampus by Ca2+-permeable AMPA receptors. Proc Natl Acad Sci U S A, 112(10), 3122–3127. 10.1073/pnas.1417498112

Kim, S., Violette, C. J., & Ziff, E. B. (2015). Reduction of increased calcineurin activity rescues impaired homeostatic synaptic plasticity in presenilin 1 M146V mutant. Neurobiol Aging, 36(12), 3239–3246. 10.1016/j.neurobiolaging.2015.09.007

Kim, S., & Ziff, E. B. (2014). Calcineurin mediates synaptic scaling via synaptic trafficking of Ca2+-permeable AMPA receptors. PLoS Biol, 12(7), e1001900. 10.1371/journal.pbio.1001900

Koren, G., Nava-Ocampo, A. A., Moretti, M. E., Sussman, R., & Nulman, I. (2006). Major malformations with valproic acid. Can Fam Physician, 52, 441–442, 444, 447. https://www.ncbi.nlm.nih.gov/pubmed/16639967

Kosik, K. S., Donahue, C. P., Israely, I., Liu, X., & Ochiishi, T. (2005). Delta-catenin at the synaptic-adherens junction. Trends Cell Biol, 15(3), 172–178. 10.1016/j.tcb.2005.01.004

Lepeta, K., Lourenco, M. V., Schweitzer, B. C., Martino Adami, P. V., Banerjee, P., Catuara-Solarz, S., … Seidenbecher, C. (2016). Synaptopathies: synaptic dysfunction in neurological disorders -A review from students to students. J Neurochem, 138(6), 785–805. 10.1111/jnc.13713

Libbey, J. E., Sweeten, T. L., McMahon, W. M., & Fujinami, R. S. (2005). Autistic disorder and viral infections. J Neurovirol, 11(1), 1–10. 10.1080/13550280590900553

Mabunga, D. F., Gonzales, E. L., Kim, J. W., Kim, K. C., & Shin, C. Y. (2015). Exploring the Validity of Valproic Acid Animal Model of Autism. Exp Neurobiol, 24(4), 285–300. 10.5607/en.2015.24.4.285

Martinez, M. C., Ochiishi, T., Majewski, M., & Kosik, K. S. (2003). Dual regulation of neuronal morphogenesis by a delta-catenin-cortactin complex and Rho. J Cell Biol, 162(1), 99–111. 10.1083/jcb.200211025

Matter, C., Pribadi, M., Liu, X., & Trachtenberg, J. T. (2009). Delta-catenin is required for the maintenance of neural structure and function in mature cortex in vivo. Neuron, 64(3), 320–327. 10.1016/j.neuron.2009.09.026

McMichael, G., Girirajan, S., Moreno-De-Luca, A., Gecz, J., Shard, C., Nguyen, L. S., … MacLennan, A. (2014). Rare copy number variation in cerebral palsy. Eur J Hum Genet, 22(1), 40–45. 10.1038/ejhg.2013.93

Medina, M., Marinescu, R. C., Overhauser, J., & Kosik, K. S. (2000). Hemizygosity of delta-catenin (CTNND2) is associated with severe mental retardation in cri-du-chat syndrome. Genomics, 63(2), 157–164. 10.1006/geno.1999.6090

Mejias, R., Adamczyk, A., Anggono, V., Niranjan, T., Thomas, G. M., Sharma, K., … Wang, T. (2011). Gain-of-function glutamate receptor interacting protein 1 variants alter GluA2 recycling and surface distribution in patients with autism. Proc Natl Acad Sci U S A, 108(12), 4920–4925. 10.1073/pnas.1102233108

Mendez-Vazquez, H., Roach, R. L., Nip, K., Chanda, S., Sathler, M. F., Garver, T., … Kim, S. (2023). The autism-associated loss of delta-catenin functions disrupts social behavior. Proc Natl Acad Sci U S A, 120(22), e2300773120. 10.1073/pnas.2300773120

Miller, D. E., Squire, A., & Bennett, J. T. (2019). A child with autism, behavioral issues, and dysmorphic features found to have a tandem duplication within CTNND2 by mate-pair sequencing. Am J Med Genet A. 10.1002/ajmg.a.61442

Moore, S. J., Turnpenny, P., Quinn, A., Glover, S., Lloyd, D. J., Montgomery, T., & Dean, J. C. (2000). A clinical study of 57 children with fetal anticonvulsant syndromes. J Med Genet, 37(7), 489–497. 10.1136/jmg.37.7.489

Moss, J. F., Oliver, C., Berg, K., Kaur, G., Jephcott, L., & Cornish, K. (2008). Prevalence of autism spectrum phenomenology in Cornelia de Lange and Cri du Chat syndromes. Am J Ment Retard, 113(4), 278–291. 10.1352/0895-8017(2008)113[278:POASPI]2.0.CO;2

Narita, M., Oyabu, A., Imura, Y., Kamada, N., Yokoyama, T., Tano, K., … Narita, N. (2010). Nonexploratory movement and behavioral alterations in a thalidomide or valproic acid-induced autism model rat. Neurosci Res, 66(1), 2–6. 10.1016/j.neures.2009.09.001

Nicolini, C., & Fahnestock, M. (2018). The valproic acid-induced rodent model of autism. Exp Neurol, 299(Pt A), 217–227. 10.1016/j.expneurol.2017.04.017

Niebuhr, E. (1978). The Cri du Chat syndrome: epidemiology, cytogenetics, and clinical features. Hum Genet, 44(3), 227–275. https://www.ncbi.nlm.nih.gov/pubmed/365706

Nivard, M. G., Mbarek, H., Hottenga, J. J., Smit, J. H., Jansen, R., Penninx, B. W., … Boomsma, D. I. (2014). Further confirmation of the association between anxiety and CTNND2: replication in humans. Genes Brain Behav, 13(2), 195–201. 10.1111/gbb.12095

Noirot, E. (1966). Ultra-sounds in young rodents. I. Changes with age in albino mice. Anim Behav, 14(4), 459–462. 10.1016/s0003-3472(66)80045-3

Ornoy, A., Weinstein-Fudim, L., & Ergaz, Z. (2015). Prenatal factors associated with autism spectrum disorder (ASD). Reprod Toxicol, 56, 155–169. 10.1016/j.reprotox.2015.05.007

Overhauser, J., Beaudet, A. L., & Wasmuth, J. J. (1986). A fine structure physical map of the short arm of chromosome 5. Am J Hum Genet, 39(5), 562–572. https://www.ncbi.nlm.nih.gov/pubmed/2878609

Peifer, M., Berg, S., & Reynolds, A. B. (1994). A repeating amino acid motif shared by proteins with diverse cellular roles. Cell, 76(5), 789–791. https://www.ncbi.nlm.nih.gov/pubmed/7907279

Portera-Cailliau, C., Pan, D. T., & Yuste, R. (2003). Activity-regulated dynamic behavior of early dendritic protrusions: evidence for different types of dendritic filopodia. The Journal of neuroscience : the official journal of the Society for Neuroscience, 23(18), 7129–7142. 10.1523/JNEUROSCI.23-18-07129.2003

Portfors, C. V. (2007). Types and functions of ultrasonic vocalizations in laboratory rats and mice. J Am Assoc Lab Anim Sci, 46(1), 28–34. https://www.ncbi.nlm.nih.gov/pubmed/17203913

Potasiewicz, A., Gzielo, K., Popik, P., & Nikiforuk, A. (2020). Effects of prenatal exposure to valproic acid or poly(I:C) on ultrasonic vocalizations in rat pups: The role of social cues. Physiol Behav, 225, 113113. 10.1016/j.physbeh.2020.113113

Rahman, M., Awosika, A. O., & Nguyen, H. (2023). Valproic Acid. In StatPearls. https://www.ncbi.nlm.nih.gov/pubmed/32644538

Ramanathan, S., Woodroffe, A., Flodman, P. L., Mays, L. Z., Hanouni, M., Modahl, C. B., … Smith, M. (2004). A case of autism with an interstitial deletion on 4q leading to hemizygosity for genes encoding for glutamine and glycine neurotransmitter receptor sub-units (AMPA 2, GLRA3, GLRB) and neuropeptide receptors NPY1R, NPY5R. BMC Med Genet, 5, 10. 10.1186/1471-2350-5-10

Rasalam, A. D., Hailey, H., Williams, J. H., Moore, S. J., Turnpenny, P. D., Lloyd, D. J., & Dean, J. C. (2005). Characteristics of fetal anticonvulsant syndrome associated autistic disorder. Dev Med Child Neurol, 47(8), 551–555. https://www.ncbi.nlm.nih.gov/pubmed/16108456

Restituito, S., Khatri, L., Ninan, I., Mathews, P. M., Liu, X., Weinberg, R. J., & Ziff, E. B. (2011). Synaptic autoregulation by metalloproteases and gamma-secretase [Research Support, N.I.H., Extramural Research Support, Non-U.S. Gov’t]. The Journal of neuroscience : the official journal of the Society for Neuroscience, 31(34), 12083–12093. 10.1523/JNEUROSCI.2513-11.2011

Robert, E., & Guibaud, P. (1982). Maternal valproic acid and congenital neural tube defects. Lancet, 2(8304), 937. https://www.ncbi.nlm.nih.gov/pubmed/6126782

Roberts, J. P., Stokoe, S. A., Sathler, M. F., Nichols, R. A., & Kim, S. (2021). Selective co-activation of alpha7- and alpha4beta2-nicotinic acetylcholine receptors reverses beta-amyloid-induced synaptic dysfunction. J Biol Chem, 100402. 10.1016/j.jbc.2021.100402

Rodier, P. M., Ingram, J. L., Tisdale, B., Nelson, S., & Romano, J. (1996). Embryological origin for autism: developmental anomalies of the cranial nerve motor nuclei. J Comp Neurol, 370(2), 247–261. 10.1002/(SICI)1096-9861(19960624)370:2<247::AID-CNE8>3.0.CO;2-2

Rossignol, D. A., Genuis, S. J., & Frye, R. E. (2014). Environmental toxicants and autism spectrum disorders: a systematic review. Transl Psychiatry, 4(2), e360. 10.1038/tp.2014.4

Roullet, F. I., Lai, J. K., & Foster, J. A. (2013). In utero exposure to valproic acid and autism--a current review of clinical and animal studies. Neurotoxicol Teratol, 36, 47–56. 10.1016/j.ntt.2013.01.004

Sandin, S., Lichtenstein, P., Kuja-Halkola, R., Hultman, C., Larsson, H., & Reichenberg, A. (2017). The Heritability of Autism Spectrum Disorder. JAMA, 318(12). doi:10.1001/jama.2017.12141

Sardina, J. M., Walters, A. R., Singh, K. E., Owen, R. X., & Kimonis, V. E. (2014). Amelioration of the typical cognitive phenotype in a patient with the 5pter deletion associated with Cri-du-chat syndrome in addition to a partial duplication of CTNND2. Am J Med Genet A, 164A(7), 1761-1764. 10.1002/ajmg.a.36494

Sathler, M. F., Doolittle, M. J., Cockrell, J. A., Nadalin, I. R., Hofmann, F., VandeWoude, S., & Kim, S. (2022). HIV and FIV glycoproteins increase cellular tau pathology via cGMP-dependent kinase II activation. J Cell Sci, 135(12). 10.1242/jcs.259764

Sathler, M. F., Khatri, L., Roberts, J. P., Schmidt, I. G., Zaytseva, A., Kubrusly, R. C. C., … Kim, S. (2021). Phosphorylation of the AMPA receptor subunit GluA1 regulates clathrin-mediated receptor internalization. J Cell Sci, 134(17). 10.1242/jcs.257972

Scattoni, M. L., Crawley, J., & Ricceri, L. (2009). Ultrasonic vocalizations: a tool for behavioural phenotyping of mouse models of neurodevelopmental disorders. Neurosci Biobehav Rev, 33(4), 508–515. 10.1016/j.neubiorev.2008.08.003

Schneider, T., & Przewlocki, R. (2005). Behavioral alterations in rats prenatally exposed to valproic acid: animal model of autism. Neuropsychopharmacology, 30(1), 80–89. 10.1038/sj.npp.1300518

Sewell, G. D. (1970). Ultrasonic communication in rodents. Nature, 227(5256), 410. 10.1038/227410a0

Shou, J., Tran, A., Snyder, N., Bleem, E., & Kim, S. (2018). Distinct Roles of GluA2-lacking AMPA Receptor Expression in Dopamine D1 or D2 Receptor Neurons in Animal Behavior. Neuroscience, 398, 102–112. 10.1016/j.neuroscience.2018.12.002

Silverman, J. B., Restituito, S., Lu, W., Lee-Edwards, L., Khatri, L., & Ziff, E. B. (2007). Synaptic anchorage of AMPA receptors by cadherins through neural plakophilin-related arm protein AMPA receptor-binding protein complexes. The Journal of neuroscience : the official journal of the Society for Neuroscience, 27(32), 8505–8516. 10.1523/JNEUROSCI.1395-07.2007

Sun, J. L., Stokoe, S. A., Roberts, J. P., Sathler, M. F., Nip, K. A., Shou, J., … Kim, S. (2019). Co-activation of selective nicotinic acetylcholine receptors is required to reverse beta amyloid-induced Ca(2+) hyperexcitation. Neurobiol Aging, 84, 166–177. 10.1016/j.neurobiolaging.2019.09.005

Sztukowski, K., Nip, K., Ostwald, P. N., Sathler, M. F., Sun, J. L., Shou, J., … Kim, S. (2018). HIV induces synaptic hyperexcitation via cGMP-dependent protein kinase II activation in the FIV infection model. PLoS Biol, 16(7), e2005315. 10.1371/journal.pbio.2005315

Takeichi, M. (1988). Cadherins: key molecules for selective cell-cell adhesion. IARC Sci Publ(92), 76-79. https://www.ncbi.nlm.nih.gov/pubmed/3235156

Taylor, B., Miller, E., Lingam, R., Andrews, N., Simmons, A., & Stowe, J. (2002). Measles, mumps, and rubella vaccination and bowel problems or developmental regression in children with autism: population study. BMJ, 324(7334), 393–396. 10.1136/bmj.324.7334.393

Thabault, M., Turpin, V., Maisterrena, A., Jaber, M., Egloff, M., & Galvan, L. (2022). Cerebellar and Striatal Implications in Autism Spectrum Disorders: From Clinical Observations to Animal Models. Int J Mol Sci, 23(4). 10.3390/ijms23042294

Tran, T. M., Sherwood, J. K., Doolittle, M. J., Sathler, M. F., Hofmann, F., Stone-Roy, L. M., & Kim, S. (2021). Loss of cGMP-dependent protein kinase II alters ultrasonic vocalizations in mice, a model for speech impairment in human microdeletion 4q21 syndrome. Neurosci Lett, 759, 136048. 10.1016/j.neulet.2021.136048

Tsuji, C., Fujisaku, T., & Tsuji, T. (2020). Oxytocin ameliorates maternal separation-induced ultrasonic vocalisation calls in mouse pups prenatally exposed to valproic acid. J Neuroendocrinol, 32(4), e12850. 10.1111/jne.12850

Tuncay, I. O., Parmalee, N. L., Khalil, R., Kaur, K., Kumar, A., Jimale, M., … Chahrour, M. H. (2022). Analysis of recent shared ancestry in a familial cohort identifies coding and noncoding autism spectrum disorder variants. NPJ Genom Med, 7(1), 13. 10.1038/s41525-022-00284-2

Turner, T. N., Sharma, K., Oh, E. C., Liu, Y. P., Collins, R. L., Sosa, M. X., … Chakravarti, A. (2015). Loss of delta-catenin function in severe autism. Nature, 520(7545), 51–56. 10.1038/nature14186

Vrijenhoek, T., Buizer-Voskamp, J. E., van der Stelt, I., Strengman, E., Genetic, R., Outcome in Psychosis, C., … Veltman, J. A. (2008). Recurrent CNVs disrupt three candidate genes in schizophrenia patients. Am J Hum Genet, 83(4), 504–510. 10.1016/j.ajhg.2008.09.011

Wang, T., Guo, H., Xiong, B., Stessman, H. A., Wu, H., Coe, B. P., … Eichler, E. E. (2016). De novo genic mutations among a Chinese autism spectrum disorder cohort. Nat Commun, 7, 13316. 10.1038/ncomms13316

Weiss, L. A., Arking, D. E., Gene Discovery Project of Johns, H., the Autism, C., Daly, M. J., & Chakravarti, A. (2009). A genome-wide linkage and association scan reveals novel loci for autism. Nature, 461(7265), 802-808. 10.1038/nature08490

Williams, G., King, J., Cunningham, M., Stephan, M., Kerr, B., & Hersh, J. H. (2001). Fetal valproate syndrome and autism: additional evidence of an association. Dev Med Child Neurol, 43(3), 202–206. https://www.ncbi.nlm.nih.gov/pubmed/11263692

Yoo, H. (2015). Genetics of Autism Spectrum Disorder: Current Status and Possible Clinical Applications. Exp Neurobiol, 24(4), 257–272. 10.5607/en.2015.24.4.257

Yuan, L., & Arikkath, J. (2017). Functional roles of p120ctn family of proteins in central neurons. Semin Cell Dev Biol, 69, 70–82. 10.1016/j.semcdb.2017.05.027

Yuan, L., Seong, E., Beuscher, J. L., & Arikkath, J. (2015). delta-Catenin Regulates Spine Architecture via Cadherin and PDZ-dependent Interactions. J Biol Chem, 290(17), 10947–10957. 10.1074/jbc.M114.632679

Zappala, C., Barrios, C. D., & Depino, A. M. (2023). Social deficits in mice prenatally exposed to valproic acid are intergenerationally inherited and rescued by social enrichment. Neurotoxicology, 97, 89–100. 10.1016/j.neuro.2023.05.009

Zaytseva, A., Bouckova, E., Wiles, M. J., Wustrau, M. H., Schmidt, I. G., Mendez-Vazquez, H., … Kim, S. (2023). Ketamine’s rapid antidepressant effects are mediated by Ca(2+)-permeable AMPA receptors. Elife, 12. 10.7554/eLife.86022

Zhang, B., Willing, M., Grange, D. K., Shinawi, M., Manwaring, L., Vineyard, M., … Cottrell, C. E. (2016). Multigenerational autosomal dominant inheritance of 5p chromosomal deletions. Am J Med Genet A, 170(3), 583–593. 10.1002/ajmg.a.37445

Zimmer, M. R., Fonseca, A. H. O., Iyilikci, O., Pra, R. D., & Dietrich, M. O. (2019). Functional Ontogeny of Hypothalamic Agrp Neurons in Neonatal Mouse Behaviors. Cell, 178(1), 44–59 e47. 10.1016/j.cell.2019.04.026

Zimprich, A., Niessing, J., Cohen, L., Garrett, L., Einicke, J., Sperling, B., … Holter, S. M. (2017). Assessing Sociability, Social Memory, and Pup Retrieval in Mice. Curr Protoc Mouse Biol, 7(7), 287–305. 10.1002/cpmo.36

