## Supplementary figures and images for "Prenatal exposure to valproic acid reduces synaptic δ-catenin levels and disrupts ultrasonic vocalization in neonates"

### Untruncated image of gels

**Fig. 1b**

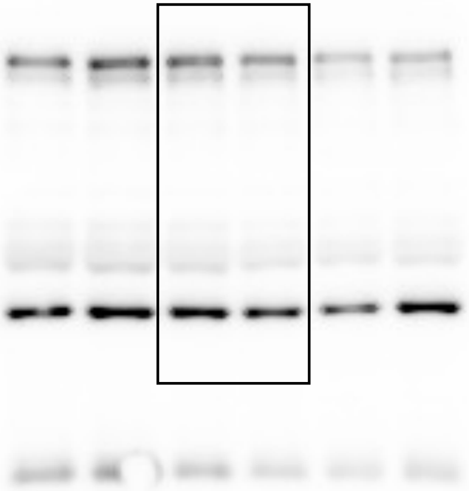


**Fig. 1c**

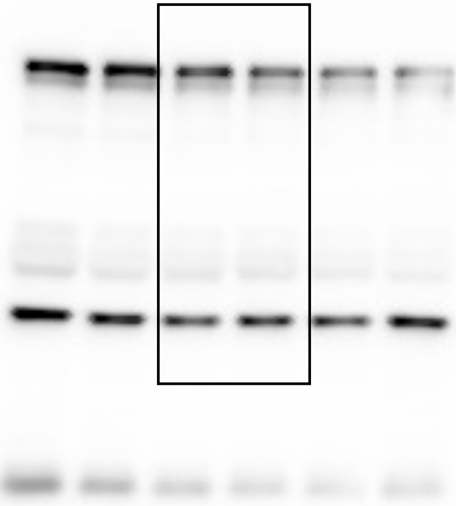


**Fig. 1a**

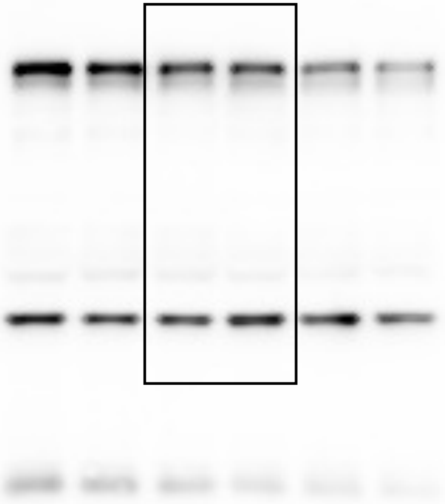

**Fig. 3 - δ-catenin**

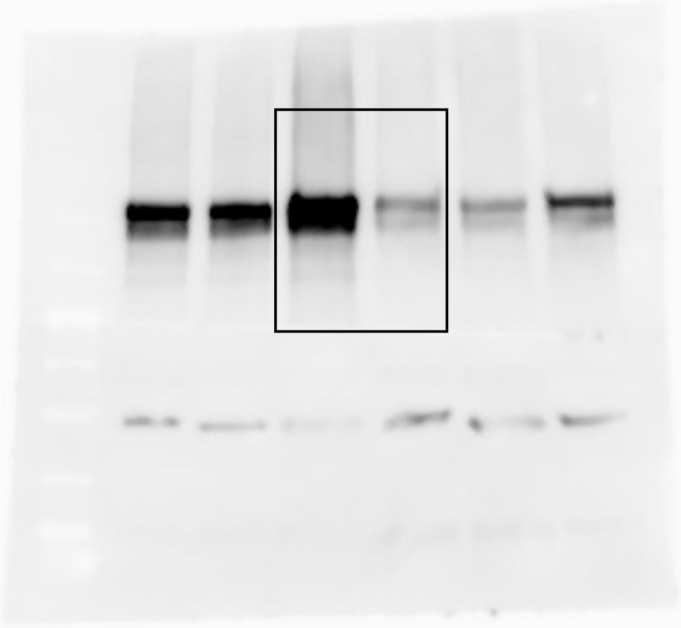

**Fig. 3 – GluA1**

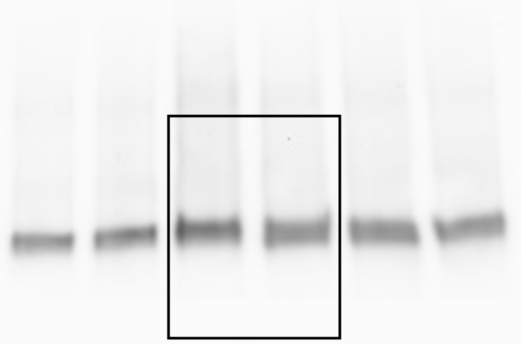

**Fig. 3 – GluA2**

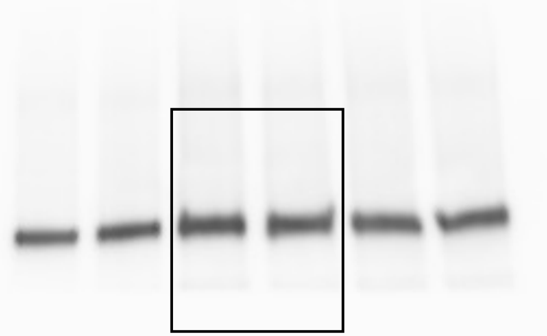

**Fig. 3 – PSD95**

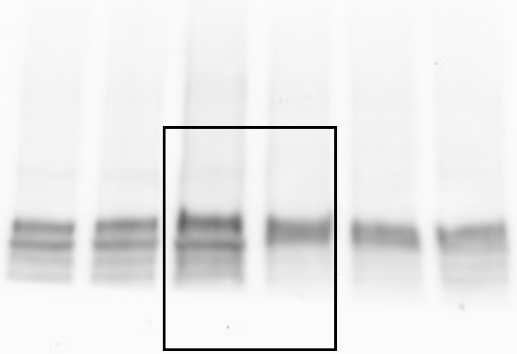

#### Fig. 3 – Actin

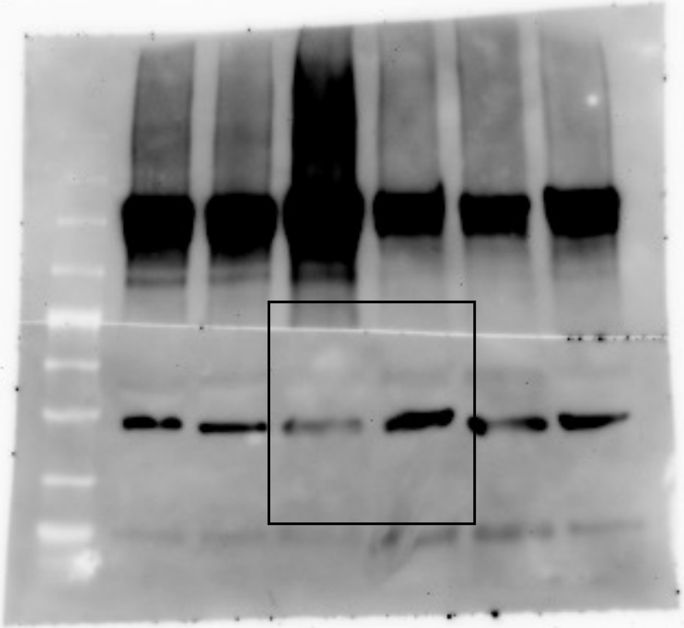
